# Antagonism of the azoles to olorofim and cross-resistance are governed by linked transcriptional networks in *Aspergillus fumigatus*

**DOI:** 10.1101/2021.11.18.469075

**Authors:** Norman van Rhijn, Sam Hemmings, Isabelle S. R. Storer, Clara Valero, Hajer Alshammri, Gustavo H. Goldman, Fabio Gsaller, Jorge Amich, Michael J Bromley

## Abstract

Aspergillosis, in its various manifestations, is a major cause of morbidity and mortality. Very few classes of antifungal drugs have been approved for clinical use to treat these diseases and resistance to the first line therapeutic class, the triazoles, is increasing. A new class of antifungals that target pyrimidine biosynthesis, the orotomides, are currently in development with the first compound in this class, olorofim in late-stage clinical trials. In this study, we identify an antagonistic action of the triazoles on the action of olorofim. We show that this antagonism is the result of an azole induced upregulation of the pyrimidine biosynthesis pathway and regulation. Intriguingly, we show that loss of function in the higher order transcription factor, HapB a member of the heterotrimeric HapB/C/E (CBC) complex or the regulator of nitrogen metabolic genes AreA, leads to cross resistance to both the azoles and olorofim indicating that factors that govern resistance are under common regulatory control. However loss of azole induced antagonism requires decoupling of the pyrimidine biosynthetic pathway in a manner independent of the action of a single transcription factor. Our study provides a first insight into antagonism between the azoles and olorofim through dysregulation of the pyrimidine and ergosterol pathway, showing complex crosstalk between these two pathways.

## Introduction

Invasive and chronic forms of aspergillosis affect over 3 million people resulting in excess of 300 thousand deaths per year [1]. Only three classes of antifungals are currently available to treat aspergillosis, with the triazoles used as first-line therapy in most centres [2]. Resistance to the azoles is rising, which is linked to use of triazole compounds in agri- and horticulture [3, 4]. It is predicted that more resistant *A. fumigatus* will be seen as azole use will be expanded to combat climate change-associated to increasing fungal crop damage [5]. The development of novel classes of antifungals will be a key component to addressing the emerging resistance problem. Fortunately, there are a number of drugs that represent novel classes of antifungal currently in development for treatment of invasive aspergillosis (IA) including ibrexafungerp, which has recently (2021) gained FDA approval for treatment of vulvovaginal candidiasis, fosmanogepix which targets GPI anchor biosynthesis and olorofim (phase 3) [6]. Olorofim (formerly known as F901318 and under development by F2G, Ltd.) is of particular interest as like fosmanogepix, it has a novel mechanism of action that has not been exploited clinically [7]. As olorofim is orally bioavailable it presents a realistic alternative to the azoles for long-term treatment of chronic and allergic infections and especially resistant infections [8]. Moreover, it could potentially be used in combination therapy strategies to supress the emergence of resistance.

Olorofim acts by inhibiting the enzyme dihydroorotate dehydrogenase (DHODH), encoded by the *pyrE* gene in *A. fumigatus*, which is a crucial enzyme within the pyrimidine biosynthesis pathway and is thus required for both DNA and RNA synthesis [7]. Structural and biochemical analysis of DHODH suggests olorofim competes with CoQ to bind to DHODH, preventing the oxidation of dihydroorotate to orotate. DHODHs are grouped into 2 classes according to sequence similarity and subcellular localisation. Both mammals and most fungi have class 2 DHODH, which is bound to the inner mitochondrial membrane [9]. The human DHODH only shares 30% protein sequence identity with the *A. fumigatus* DHODH and olorofim has also been demonstrated to be >2,200-fold more potent against the *A. fumigatus* enzyme [7]. Inhibition of the pyrimidine biosynthesis pathway by olorofim prevents the germination of *A. fumigatus* conidia and causes hyphae to undergo morphological changes [10]. Prolonged exposure of germlings and vegetative hyphae to olorofim also causes extensive isotropic expansion that is then followed by cell lysis [11].

Olorofim has an effect on a wide range of fungi and has been shown to be effective against *Coccidioides immitis*, *Scedosporium* spp., *Madurella mycetomatis*, *Lomentospora prolificans* and several *Aspergillus* species [12–18]. However, olorofim has a reduced activity against *Fusarium solani* species complex and *Fusarium dimerum* and is inactive against Mucorales [19]. Olorofim is also effective against triazole resistant *A. fumigatus* isolates and cryptic *Aspergillus* species [20, 21]. In several murine models of aspergillosis, olorofim treatment significantly reduced fungal burden and mortality [15].. Reassuringly, a recent study suggests that levels of resistance to olorofim in a collection of clinical isolates of *A. fumigatus* is low. Only 1 of 976 clinical isolates exhibited pre-existing olorofim resistance caused by a single SNP in the *pyrE* gene [22].

In this study, we identify a concerning antagonistic effect of the triazoles on the action of olorofim in *A. fumigatus*. We show that this antagonistic effect is governed by an azole induced upregulation of the pyrimidine biosynthetic pathway. However, it does not appear to be regulated by the action of a single transcription factor. Through screening the COFUN *A. fumigatus* transcription factor null mutant library we identify four transcription factors that regulate susceptibility to olorofim [23]. Existing published literature, and our phenotypic and transcriptomic data revealed these transcription factors regulate genes involved in processes immediately upstream of the pyrimidine biosynthesis pathway. Notably two transcription factor null mutants, Δ*hapB* and Δ*areA,* have elevated MICs to olorofim and are resistant to the azole class of antifungals, highlighting potential routes to cross resistance.

## Materials and Methods

### Fungal strains

Conidia of *Aspergillus fumigatus* MFIG001 (a derivative of CEA10) and transcription factor null mutants [23, 24] were prepared by inoculating strains in vented 25cm^2^ tissue culture flasks with Sabouraud Dextrose agar (Oxoid, Hampshire, England) and incubating at 37°C for 48 hours. Spores were harvested in PBS + 0.01% Tween-20 by filtration through Miracloth. Spores were counted using a haemocytometer (Marienfeld Superior, Baden-Württemberg, Germany).

### Olorofim MIC screening

Olorofim was a kind gift of F2G Ltd. The Minimum Inhibitory Concentration (MIC) of olorofim against *A. fumigatus* was assessed using the European Committee for Antimicrobial Susceptibility Testing (EUCAST) methodology [19, 25]. Briefly, 2×10^⁴^ spores/mL (in 100 µl) were added to a CytoOne^®^ 96-well plate (StarLab, Brussels, Belgium) containing 1xRPMI-1640 medium (Sigma Aldrich, St. Louis, MO), 165 mM MOPS buffer (pH 7.0), 2% glucose, with olorofim 2-fold dilution series ranging from 0.1 μg/L to 0.25 mg/L and a drug free control (n = 4). Additionally, a serial dilution of olorofim containing 10 mM uracil and uridine was performed. 96-well plates were incubated at 37°C for 48 hours. The MIC was determined as the minimum drug concentration at which no germination was observed. Optical density was measured at 600 nm using a Synergy™ HTX Multi-Mode Microplate Reader (BioTek, Winooski, VT). In keeping with research laboratory based definitions, but in contract to definitions used clinically, we define *in vitro* resistance as a strain that is less susceptible to drug than the parental isolate [26].

### Olorofim sensitivity screening of the A. fumigatus transcription factor null mutant library

2×10^⁴^ spores/mL from each of the 484 members of the transcription knockout library were added to 1x RPMI-1640 medium, 165 mM MOPS buffer (pH 7.0), 2% glucose in each well of a CytoOne^®^ 96-well plate with 0.002 mg/L olorofim (n = 4). Plates were incubated at 37°C for 48 hours. Fitness was calculated by dividing the optical density of respective null mutants to the MFIG001 control. Relative fitness in olorofim was calculated by dividing fitness in olorofim with general growth fitness of the transcription factor null mutants using the same microculture conditions in 1x RPMI-1640 medium, 165 mM MOPS buffer (pH 7.0), 2% glucose without olorofim (n = 4). Optical density was measured at 600 nm on a Synergy™ HTX Multi-Mode Microplate Reader (BioTek, Winooski, VT).

### RNA-extraction

1×10^6^ spores/mL of *A. fumigatus* MFIG001, Δ*AFUB_056620* and Δ*AFUB_030440* were inoculated into 50 mL of *Aspergillus* complete media (ACM) [27] and incubated for 18 hours at 37°C in a rotary shaker (180 rpm). Mycelia were harvested using filtration through Miracloth (Merck Millipore) and washed in 1x RPMI-1640 medium. Approximately 1g of mycelia was added to shake flasks containing 50 mL RPMI-1640 medium, 165 mM MOPS buffer (pH 7.0), 2% glucose and incubated for 1 hour at 37°C in a rotary shaker (180rpm) in the presence or absence of 0.062 mg/L olorofim (n = 3), or in the presence or absence of 0.25 mg/L, 0.5 mg/L, 1 mg/L or 2 mg/L itraconazole (n=3) incubated for 4 hours. Mycelia was filtered through Miracloth and snapfrozen using liquid nitrogen and kept at −80°C until required.

To extract RNA, 1 mL of TRIzol reagent (Sigma Aldrich) and 710-1180 μm acid washed glass beads (Sigma Aldrich) were added to frozen mycelia and placed in a TissueLyser II^®^ (Qiagen, Hilden, Germany) for 3 minutes at 30 Hz. The solution was centrifuged (12,000 rpm) for 1 minute at 4**°**C. The aqueous phase was added to 200 μL of chloroform and centrifuged (12,000 rpm) for 10 minutes at room temperature. The supernatant was added to 0.2 M sodium citrate, 0.3 M sodium chloride and 25% (v/v) isopropanol and left at room temperatures for 10 minutes. This solution was centrifuged (12,000 rpm) for 15 minutes at 4**°**C. The supernatant was removed; the pellet was washed in 70% (v/v) ethanol and resuspended in RNase free water (Thermo Fisher Scientific, Waltham, MA). RNA samples were treated with RQ1 RNase-Free DNase (Promega, Madison, WI) and purified using a RNeasy Mini Kit (Qiagen). RNA quality and quantity were assessed using gel electrophoresis and using a NanoDrop^TM^ 2000/2000c Spectrophotometer (Thermo Fisher Scientific). All RNA extractions were carried out in triplicate.

### Transcriptomic Analysis

RNA sequencing was carried out by the Genomic Technologies Core Facility (GTCF) at The University of Manchester. Sequencing libraries were prepared from mRNA using TruSeq^®^ Stranded mRNA assay (Illumina, San Diego, CA). Samples were sequenced on a single lane on an Illumina HiSeq2500 (Illumina). Low-quality reads of resulting fastq files were removed using FastQC and trimmed using Trimmomatic (Quality >20, Sliding window average of 4 bases) [28]. Bowtie was used to align libraries to the *A. fumigatus* A1163 genome assembly GCA_000150145.1 with gene annotation from CADRE/Ensembl Fungi v24 [29]. Differential expression analysis of was performed using DESeq2 [30].

Functional category and gene ontology enrichment analysis was carried out using FungiFun2 2.2.8, converting genes to Af293 gene names to allow using the KEGG option [31]. Genes that showed over 2-fold in differential expression and Benjamin-Hochberg FDR <0.01 underwent enrichment analysis. StringsDB analysis was performed by only including genes with at least two connections.

### Phenotypic analysis

For colony images, 500 spores per isolate were spotted onto solid ACM or Aspergillus Minimal Media (AMM) and left to dry. Plates were incubated at 37°C for 72 hours and imaged. Growth on solid AMM supplemented with different nitrogen sources (50 mM ammonium tartrate, 10 mM sodium nitrate, 10 mM L-glutamine, 10 mM urea or 10 mM L-proline) were assessed by spotting 500 spores from each isolate (n = 3). Plates were incubated at 37°C for 72 hours. MICs were determined using the same supplementation as the phenotypic test with a serial dilution of olorofim (ranging from 0.1 μg/L to 0.25 mg/L). 96-well plates were incubated for 48 hours at 37°C and growth was determined by microscopic evaluation.

### Checkerboard assays

For assessing drug combination efficacies of itraconazole and olorofim against *A. fumigatus*, we used a checkerboard assay similar to EUCAST MIC testing described above. Twofold serial dilutions of itraconazole were prepared across the X-axis and olorofim serial dilutions across the Y-axis. The MIC was determined by microscopy by visually assessing the well containing the lowest drug concentration with non-germinated spores. The fractional inhibitory concentration index (FICI) was calculated as the MIC in combination divided by the MIC of individual drugs [32].

### Generation of TetOFF mutants

The tetOFF cassette was amplified from pSK606 [33] containing 50 bp homology arms targeted to the promoter of each target gene (Supplementary Table 1). These PCR products were used as repair template for CRISPR-Cas9 mediated transformation [34] using corresponding crRNA for each gene (Supplementary Table 1). Transformants were selected using pyrithiamine (concentration) containing AMM+1% sorbitol plates, purified twice and validated by PCR.

### Disk assays

4 ×10^4^ conidia of the relevant *A. fumigatus* strain were evenly distributed on solidified 1xRPMI 1640 (Sigma), 165 mM MOPS buffer (pH 7.0), 2% glucose. One 6 mm antibiotic assay disk (Whatman) was placed on the middle of the plate or two disks at a fixed distance, and 10 µL of voriconazole (800 mg/L), olorofim (500 mg/L), manogepix (250 mg/L) or H_2_O_2_ (30%) were added to each of them. The plates were incubated at 37°C for 48 hours and imaged. Antagonism was measured as the area within the halo when two antifungals are combined showing fungal growth. Measurements were done using FIJI.

### Data availability

RNA-seq data is available from ArrayExpress as experiment: E-MTAB-10590. The differential expression output from DESeq2 is included as Supplementary Data 1. (reviewer access: Reviewer_E-MTAB-10590 Password: pptwwqmj). Itraconazole RNA-seq is available from GEO: PRJNA861909

## Results

### The azoles are antagonistic to the action of olorofim in a manner consistent with azole mediated upregulation of the pyrimidine biosynthetic pathway

In order to standardise assays throughout our experiments, the Minimum Inhibitory Concentration (MIC) of olorofim against *Aspergillus fumigatus* MFIG001 was determined. The MIC was defined as the minimum concentration of olorofim at which no germination from *Aspergillus* spores was observed. Microscopic evaluation revealed the MIC of olorofim to be 0.06 mg/L for *A. fumigatus* MFIG001, consistent with previous findings of other *A. fumigatus* isolates [20]. The effect of olorofim on growth of *A. fumigatus* was further evaluated by measuring optical density of the plates used to determine the MIC (**Figure 1a**). The maximal growth observed (OD_600_ = 0.39) and minimal growth observed (OD_600_ = 0.04) was separated by a 64-fold difference in drug concentration, showing the effect of olorofim is progressive over a long range of concentrations until achieving total growth inhibition. This is in stark contrast to the inhibitory effects of the azoles on *A. fumigatus* where the difference between maximal and minimal growth typically occurs over a drug concentration not exceeding 8-fold (**Supplemental Figure 1**). As this range is so broad, we consider it useful to measure the concentration at which growth is inhibited by 50% (herein referred to as IC50 [35] to distinguish from MIC50 which is an MIC determination made of populations). For MFIG001, the IC50 for olorofim is 0.0057 mg/L whereas for itraconazole its 0.21 mg/L. As olorofim inhibits pyrimidine biosynthesis, it would be expected that the action of the drug would be fully reversed by supplementing the media with an excess of exogenous pyrimidines [7]. To confirm growth inhibition was due to directly targeting the pyrimidine biosynthesis pathway, the MIC was determined with the addition of 10 mM uridine and 10 mM uracil (**Figure 1b**). Under these conditions there was no observed reduction in *A. fumigatus* growth, and at all olorofim concentrations the median OD_600_ did not fall below control levels indicating that there are no significant off target effects of this drug.

**Figure 1:**
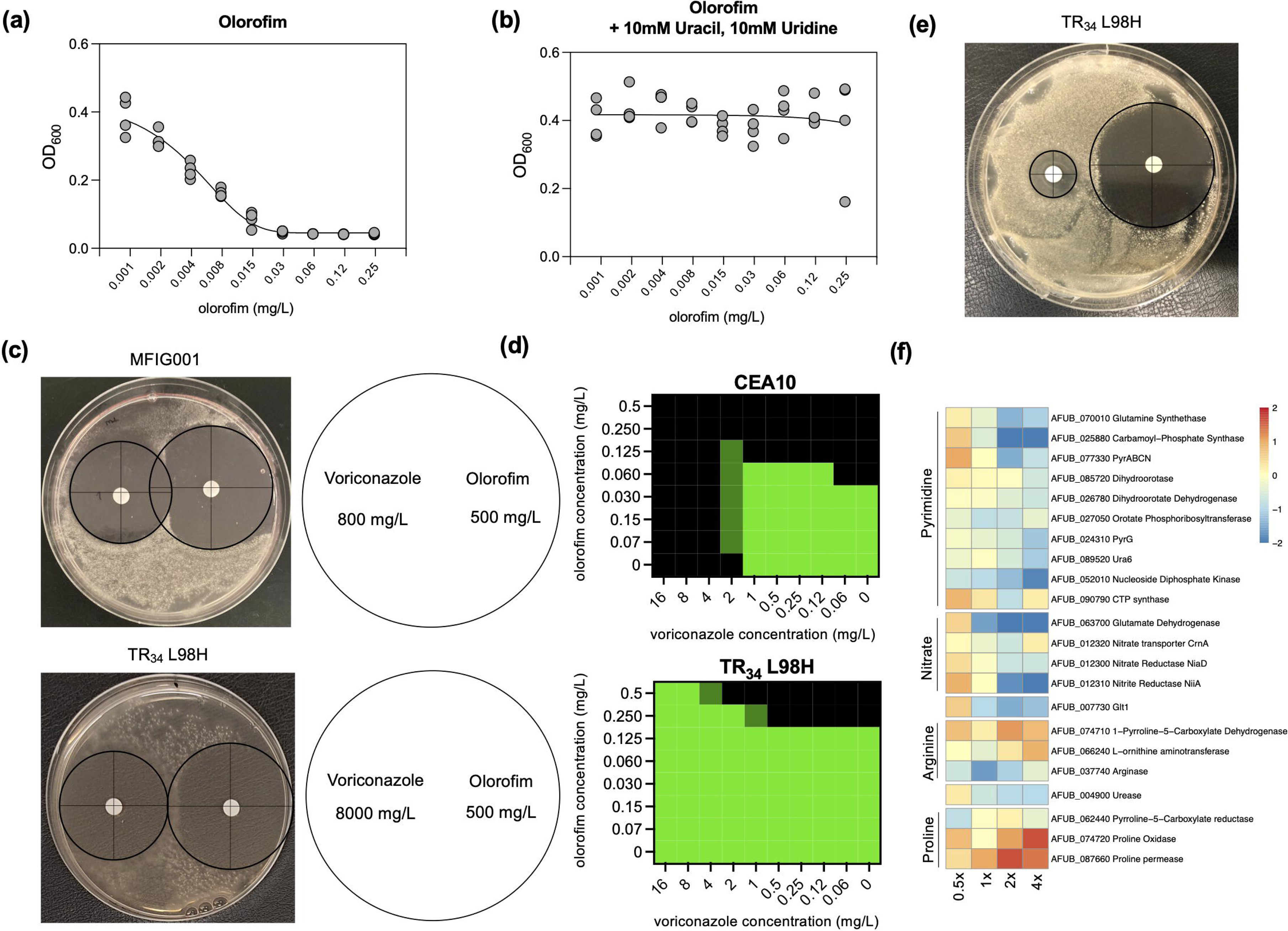
Antagonism of the azoles to olorofim. (a) Broth dilution assay of olorofim on *A. fumigatus* MFIG001 to olorofim, following EUCAST methodology and measured by OD_600_ (n=4). (b) Addition of 10mM uracil and 10mM uridine reverses the action of olorofim on *A. fumigatus* MFIG001 (n=3). (c) Antagonism on a solid RPMI-1640 plate inoculated with *A. fumigatus* isolates. Voriconazole (800 mg/L) is inoculated on the disk on the left, olorofim (500 mg/L) on the disk on the right. The disk assay for TR34 L98H contained 8000 mg/L voriconazole to obtain a halo in equal size to MFIG001. (d) Checkerboard assay (n=3) for CEA10 and the azole-resistant TR34 L98H isolate to voriconazole and olorofim. Growth is normalised to RPM-1640 without any antifungal drug. Green equals full growth, black no observed growth. (e) Disk assay on solid RPMI-1640 at 800 mg/L voriconazole and 500 mg/L olorofim for the TR34 L98H strain. (f) Dose response RNA-seq upon itraconazole exposure (0.5x MIC – 4x MIC). Expression of genes of the pyrimidine pathway and upstream pathways are differentially upregulated only in sub-MIC concentrations of itraconazole

Resistance to the clinical azoles has become a global problem that is being addressed in multiple centres by using combination therapy with either an echinocandin or amphotericin B. If approved for use, olorofim may be used in the same way. We therefore investigated the potential interaction in activity between the triazoles; voriconazole and itraconazole, and olorofim against CEA10, MFIG001 and a TR34 L98H azole-resistant isolate generated in the MFIG001 background [36]. To our surprise given the distinct mechanisms of action of the orotomides and the azoles, we observed a clear uni-directional antagonism by the azoles on olorofim in both liquid and solid media **(Figure 1c and d**). Interestingly we did not see the same antagonism between olorofim and manogepix, another late stage antifungal compound (**Supplemental Figure 2**). The antagonism of the azoles to olorofim was also observed under non-growth inhibitory concentrations of voriconazole for the TR34 L98H azole-resistant isolate, showing that this antagonistic response is independent of the azole antifungal activity (**Figure 1e**).

To gain an understanding of the potential mechanisms driving this antagonism, we evaluated transcriptomic data for *A. fumigatus* MFIG001 exposed to increasing concentrations of itraconazole (**Figure 1f**). As expected, the ergosterol biosynthetic pathway was differentially regulated throughout itraconazole concentrations. At sub-MIC levels of itraconazole we observed a significant upregulation of genes in the pyrimidine biosynthetic pathway and those pathways that generate its precursors **(Supplemental Data 1**). Most strikingly, the nitrate assimilation pathway, *glt1* and the first three steps in the pyrimidine pathway (encoded by *glnA*-AFUB_070010, *pyrD*-AFUB_085720 and *pyrABCN-AFUB_077330* and its orthologues *AFUB_025880* and *AFUB_054340*) were upregulated in sub-MIC levels of itraconazole (**Figure 1e**); interestingly many of these genes were downregulated in supra-MIC concentrations of itraconazole suggesting metabolic arrest [37]. This led us to hypothesise that both the pyrimidine pathway and ergosterol biosynthesis pathways are potentially co-regulated.

### Deletion of HapB, AreA, DevR and AcdX changes olorofim susceptibility

As we observed antagonism between the azoles and olorofim, and co-regulation of those pathways upon azole exposure, we hypothesised that both pathways may be co-regulated by the same transcription factors. To assess this co-regulation and identify novel transcriptional regulators associated with differential olorofim susceptibility and azole antagonism, the COFUN transcription factor knockout (TFKO) library was screened against olorofim at a concentration that reduces growth of the isogenic wildtype isolate (MFIG001) by about 20% (0.002 mg/L). At this concentration we were able to identify strains that have the potential to be resistant or hypersensitive (**Figure 2a**) while utilising resource limiting levels of drug.

**Figure 2:**
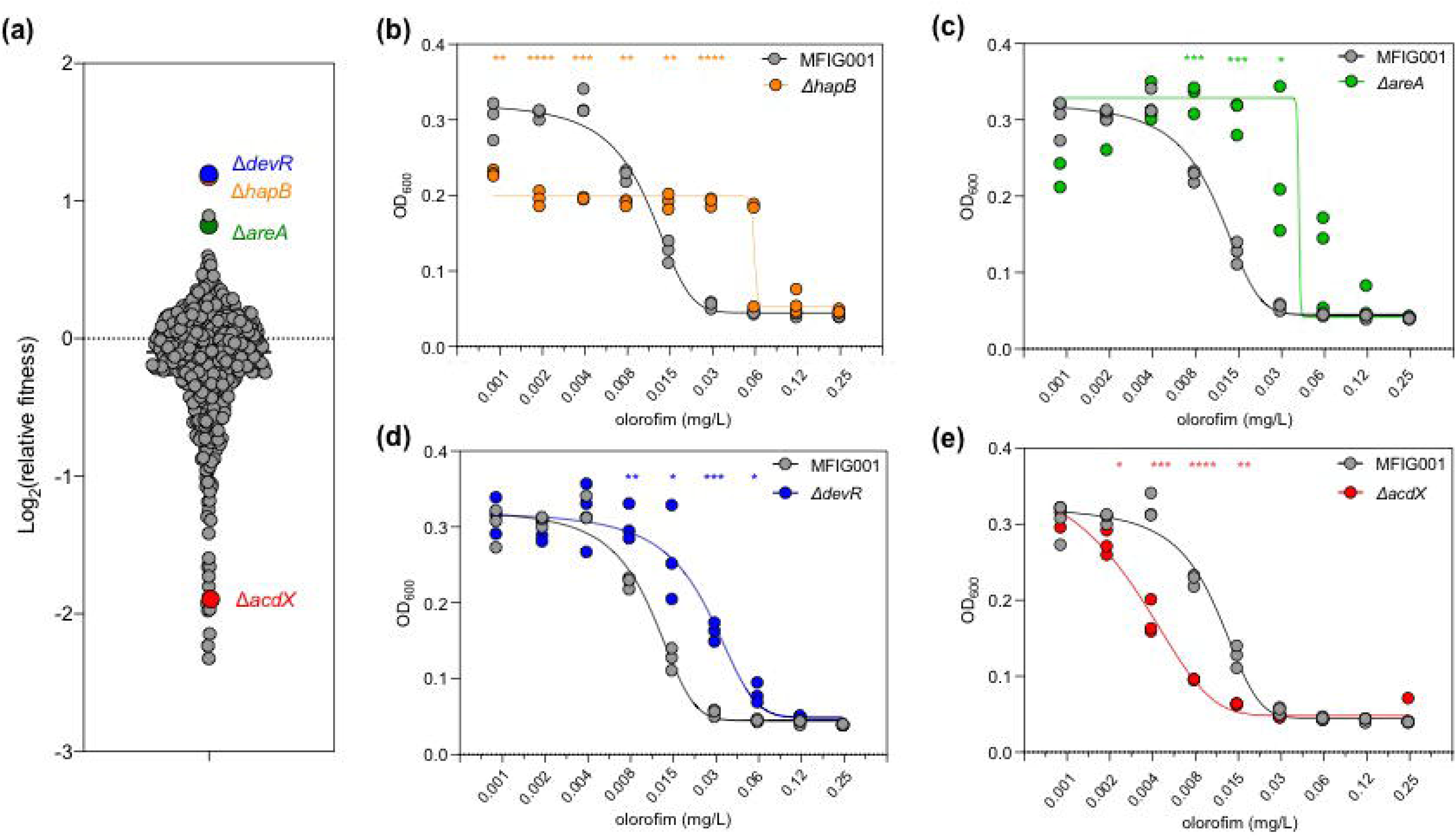
Olorofim susceptibility screening of the COFUN transcription factor knockout library. (a) Relative fitness of each individual strain was assessed by normalising to fitness in non-drug condition (n=3). TF null mutants that are of particular interest are highlighted. (b-e) Broth dilution assay of olorofim on the TF null mutants, (b) for Δ*hapB*, (c) for Δ*areA*, (d) for Δ*devR*, (e) for Δ*acdX*, as determined by OD_600_ (n=3). Statistical difference was assessed by Two-way ANOVA with Sidaks multiple comparison test (* p<0.05, ** p<0.01, *** p<0.001, *** p<0.001).

Three transcription factor null mutants (Δ*areA,* Δ*hapB* and Δ*devR*) showed reproducible increased relative fitness in the presence of olorofim and elevated MICs compared to MFIG001 (**Figure 2b, c and d**). Remarkably, two of these mutants (Δ*areA* and Δ*hapB*) are also resistant to the azole class of antifungals [23]. Loss of AreA, a transcription factor that has a global role in activating expression of genes involved in nitrogen acquisition and processing [38] or loss of HapB, which along with HapC and HapE comprise the CCAAT Binding Complex (CBC) [39] resulted in a 2-fold increase in MIC to olorofim when compared to the isotype control MFIG001; IC50 values for these strains were 0.04 mg/L (4-fold increase) and 0.07 mg/L (8-fold increase), respectively (**Figures 2b and 2c**). This simultaneous decrease in azole and olorofim susceptibility suggests suggesting a higher level regulatory link between ergosterol biosynthesis and pyrimidine biosynthesis. DevR is a bHLH transcription factor involved in sporulation and melanin biosynthesis [40]. The Δ*devR* mutant showed a significant reduction in susceptibility to olorofim at concentrations ranging from 0.008 mg/L to 0.06 mg/L (MIC) and had an IC50 of 0.025 mg/L (**Figure 2d**). Although the MIC for this strain increased to >0.125 mg/L most spores did not germinate at this concentration.

One isolate (ΔAFUB_056620, Δ*acdX*) showed a reproducible significant increase in sensitivity to olorofim and had an MIC of 0.03 mg/L and a IC50 of 0.006 mg/L, 2-fold lower than *A. fumigatus* MFIG001 (**Figure 2e**). The *acdX* gene encodes a 612 amino acid transcription factor that contains six WD40 repeat units but no other functional domains, as shown by a SMART domain search. A reciprocal BLAST of the AFUB_056620 protein sequence found a match to the *Saccharomyces cerevisiae* transcription factor Spt8. However, the proteins only share 44% identity of the entire protein sequence. In *S. cerevisiae*, Spt8 forms part of the SAGA (Spt-Ada-Gcn5-acetyltransferase) complex [41] which is known to act as a transcriptional activator under several stress conditions. While the orthologue of AcdX in other fungi generally contain six WD40 domains, in species such as *N. crassa* and *A. terreus* only five domains are present, however the significance of this is unclear. In *A. nidulans* AcdX has been described to be functional in the SAGA complex and is involved in repressing genes in in acetate metabolism and has a regulatory role in the proline metabolic pathway [42].

### Transcription factor mutants with altered suseptibility to olorofim have defects in nitrogen assimilation

Further phenotypic analysis of the null mutants with differential susceptibility to olorofim revealed that all had differential growth on *Aspergillus* Complete Medium (ACM) (**Figure 3a and b**) and *Aspergillus* Minimal Medium (AMM), which contains ammonium tartrate as nitrogen source (**Figure 3a and c**). The *hapB, devR, areA* and *acdX* null mutants showed a reduction of radial growth on ACM of 28%, 22%, 12% and 24% respectively when compared to the isotype control (p<0.05). On AMM, the *hapB* mutant showed increase radial growth (58%) however; colony growth was more diffuse than the isotype strain (**Figure 3a and c**). As olorofim inhibits DHODH, which acts within the pyrimidine biosynthetic pathway we hypothesised that these growth defects could be reflecting an alteration in the abundance of precursors of this pathway. Substitution of ammonium tartrate to nitrate did not rescue any of the growth defects of the transcription factor null mutants, and even exacerbated them in Δ*hapB* and Δ*areA* (**Figure 3d, Supplemental Figure 3**). Glutamine supplementation rescued the growth rate defects of Δ*acdX* and Δ*areA* although significant growth defects were still present even after supplementation (**Figure 3e**). Similarly, urea almost completely rescued Δ*acdX* and proline fully rescued Δ*hapB*, Δ*devR* and Δ*acdX* (**Figure 3f and g**). Taken together, these results show that these transcription factor null mutants have defects in nitrogen utilisation that, given its connection with the pyrimidine pathway, could be linked to olorofim susceptibility.

**Figure 3:**
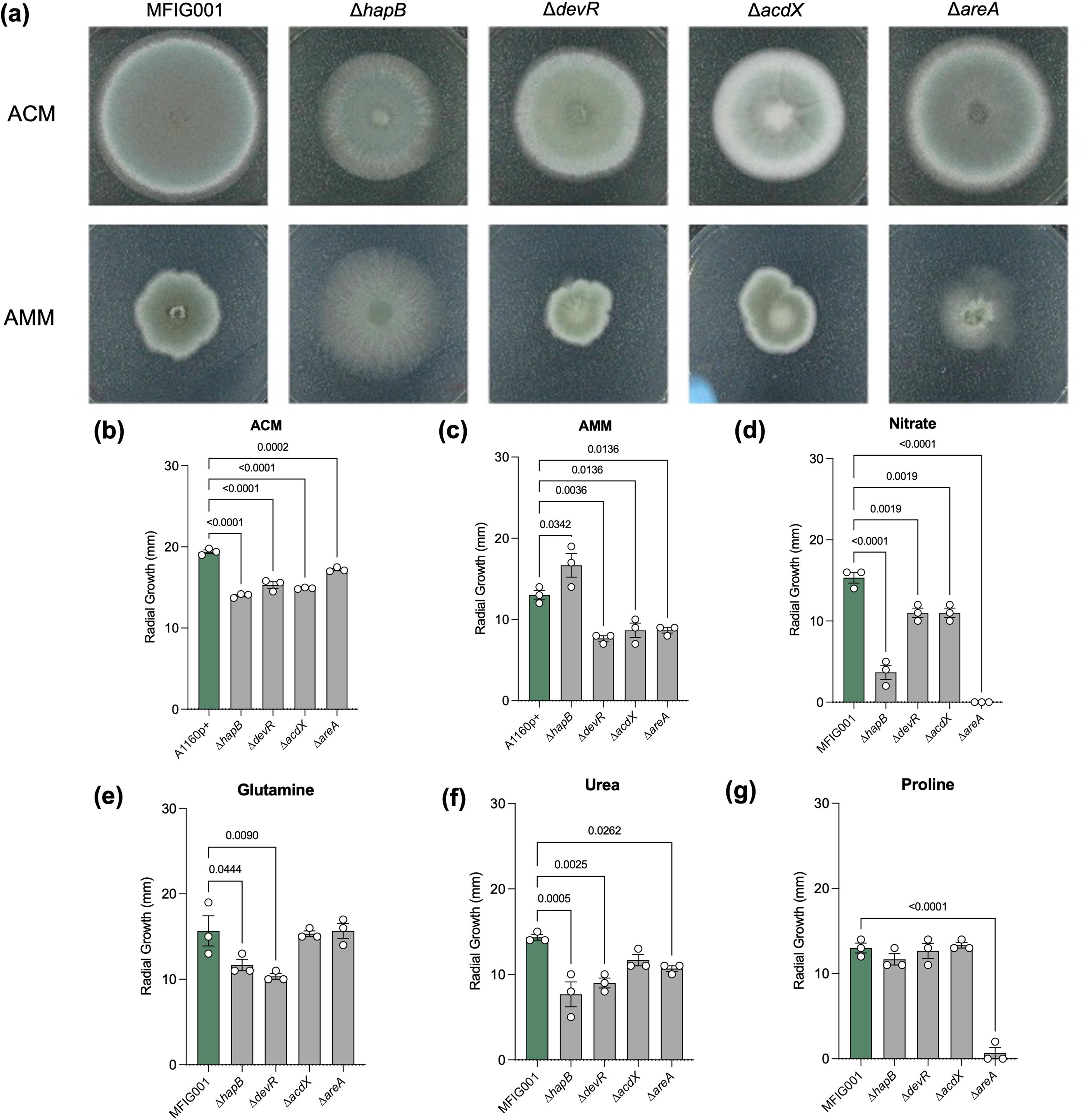
Phenotypic evaluation of TF null mutants. (a) 500 spores of TF null mutants and MFIG001 were spotted on Aspergillus Complete Medium and Aspergillus Minimal Medium and incubated for 48 hours at 37° Celsius. (b-c) Radial growth of TF null mutants and MFIG001 on ACM (b) and AMM (c), after 72 hours at 37° Celsius (n=3) (d-g) TF null mutants spotted on AMM supplemented with 10 mM sodium nitrate (d), 10 mM L-glutamine (e), 10 mM urea (f) or 10 mM L-proline (g) (n=3) Statistical difference was assessed by two-way ANOVA with Dunn’s correction (p-values < 0.05 are shown).

### Changes in susceptibility to olorofim in ΔdevR and ΔacdX mutants are caused by opposing regulation of pathways preceding pyrimidine biosynthesis

To facilitate our understanding of how these transcritpon factors are functioning to alter olorofim sensitivity we performed whole transcriptome analysis. Upon olorofim exposure (1x MIC) for 1 hour, a modest 41 genes and 185 genes were up- and downregulated Log2 fold > 1 (**Figure 4a**) in our isotype-type strain, respectively. Our expectation was that several genes in the immediate pyrimidine biosynthesis pathway would be upregulated but only the gene encoding the multifunctional carbamoyl-phosphate synthase/aspartate carbamoyltransferase (PyrABCN, AFUB_077330) enzyme, which is upstream of DHODH and converts carbamoyl-P to N-carbamoyl-L-aspartate, was upregulated by Log2 fold >1 (Supplemental Data 1). Instead, genes associated with pathways that synthesise precursors of the pyrimidine biosynthetic pathway were identified including oxaloacetate metabolism and glutamate biosynthesis (**Figure 4b, Figure 4c and Supplemental Data 1**). Genes associated with tyrosine metabolism; secondary metabolite biosynthesis, glycolysis/gluconeogenesis and valine, leucine and isoleucine degradation were enriched among downregulated genes (**Figure 4b**). A STRINGS analysis of differentially regulated genes showed an interconnected network of genes involved in ergosterol biosynthesis, the TCA cycle and nitrogen metabolism (**Figure 4c**).

**Figure 4:**
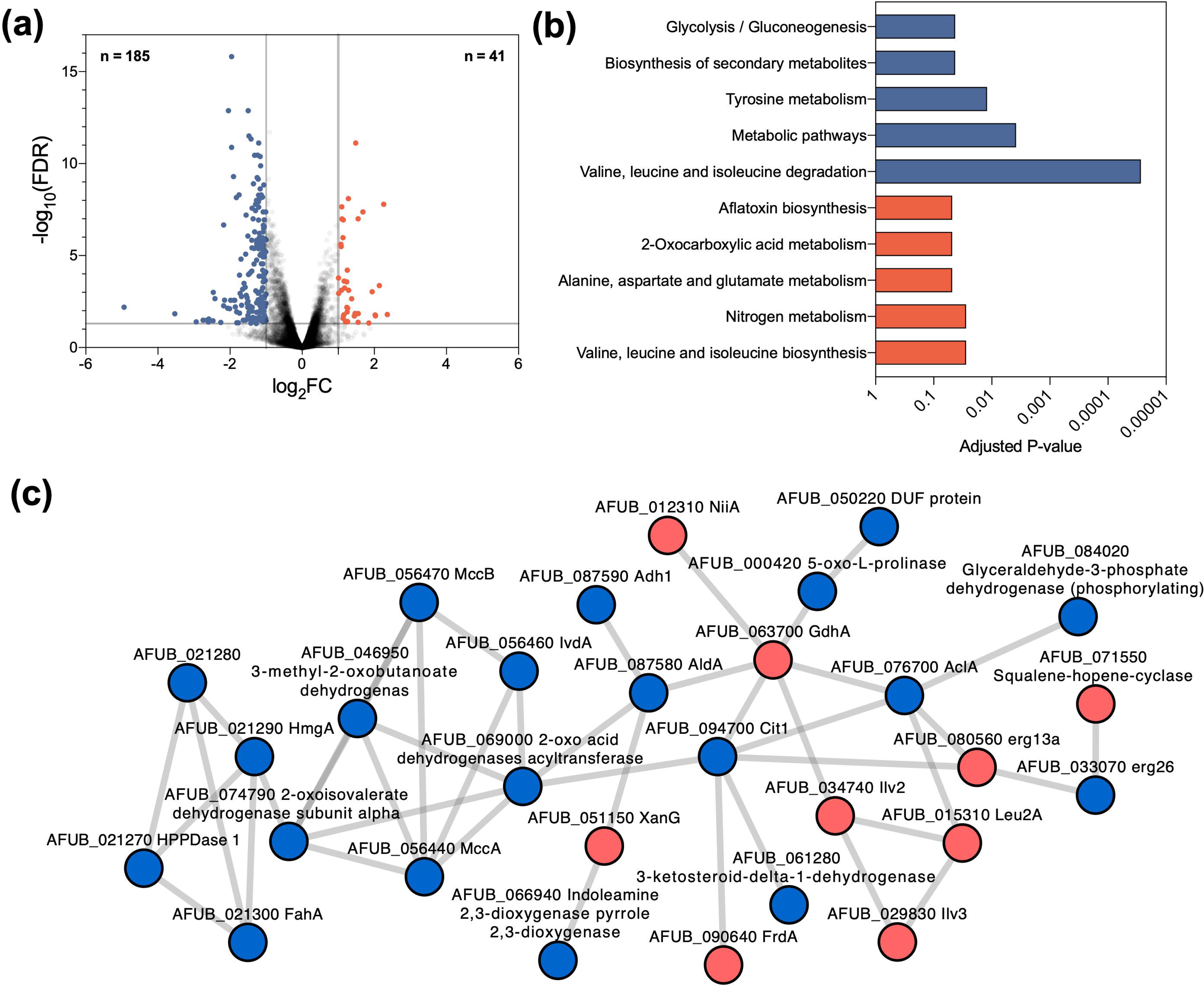
Transcriptomics of MFIG001 to olorofim. (a) Volcano plot of RNA-seq of *A. fumigatus* MFIG001 exposed to olorofim. 185 genes (blue dots) and 41 genes (red dots) were considered downregulated and upregulated, respectively (>2-fold differentially regulated, p<0.05). (b) KEGG pathways that are enriched within differentially regulated genes, blue categories are associated with downregulated genes, red with upregulated genes. (c) Interactions of proteins involved in response to olorofim as determined by StringsDB. Proteins derived from upregulated transcripts are in red, downregulated in blue.

In order to characterise the basis of differential olorofim susceptibility in the Δ*devR* and Δ*acdX* mutants the transcriptomes of these two mutants were compared to the wild-type (**Supplemental Data 2**). In the absence of olorofim 510 and 137 genes were respectively downregulated and upregulated in the Δ*devR* isolate while 212 were downregulated and 194 upregulated upon olorofim exposure. In the absence of olorofim, notable enriched functional categories included downregulation of genes involved in tyrosine metabolism and an upregulation of genes involved in the biosynthesis of branched chain amino acids and metabolism of arginine and proline, the latter of which was also seen under olorofim exposure (**Figure 5a**). A detailed pathway analysis under olorofim challenge of genes involved in the conversion of metabolites towards L-glutamate and through to orotate revealed that proline uptake and degradation were upregulated in the *devR* null mutant (**Figure 5b and d**). Other pathways that contribute to orotate precursors were also significantly upregulated, notably the nitrate assimilation pathway (NAP [*crnA, niaD, niiA*]), and glutamate, glutamine and carbomyl-P synthesis. Pathways that compete with orotidine biosynthesis for L-glutamate were not differentially regulated in any of the assessed mutants (**Supplemental Data 2).** Our transcriptional data therefore suggests that nitrogen metabolism is probably altered in this strain in ways that favor the generation of precursors for orotate biosynthesis and hence could explain the reduced sensitivity of *devR* null mutant to olorofim. The transcriptome of the olorofim hypersensitive Δ*acdX* mutant also revealed that proline and arginine metabolism were upregulated compared to the wild-type but genes involved in the NAP and glutamate, glutamine and carbomyl-P synthesis pathways were downregulated suggesting that AcdX and DevR have directly opposing functions on these linked pathways (**Figure 5c, Figure 5d**) and providing further evidence to suggest that regulation of these pathways is important for olorofim sensitivity.

**Figure 5:**
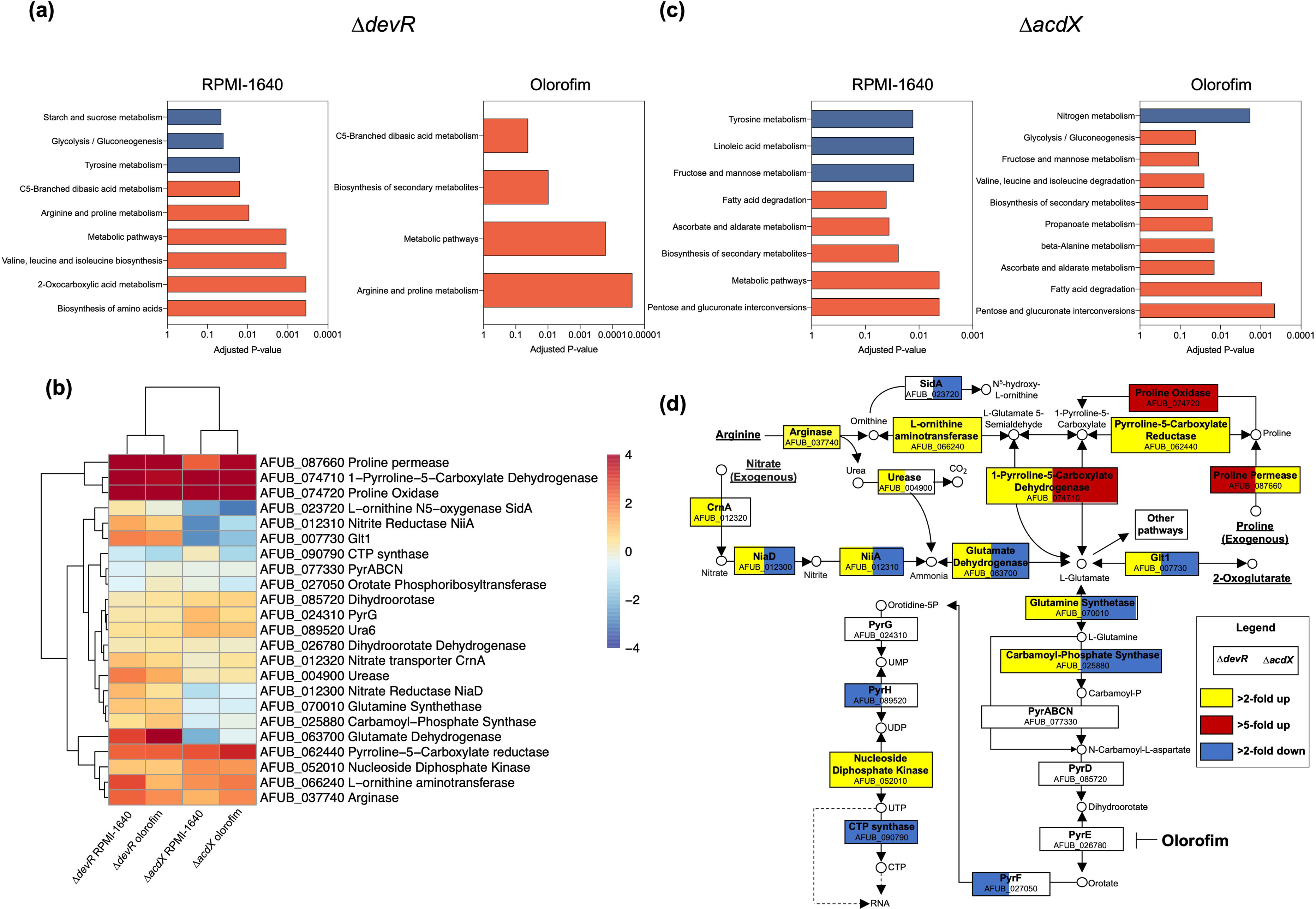
RNA-seq analysis of Δ*devR* and Δ*acdX* exposed to olorofim. (a) KEGG pathways enriched for down-(blue) or upregulated (red) genes in RPMI-1640 or upon olorofim exposure for Δ*devR* compared to *A. fumigatus* A1160p+. (b) Heatmap of genes involved in the pyrimidine pathway and component upstream of this pathway. (c) KEGG pathways enriched for down-(blue) or upregulated (red) genes in RPMI-1640 or upon olorofim exposure for Δ*acdX* compared to *A. fumigatus* MFIG001. (d) Detailed analysis of genes involved in pathways upstream of and including the pyrimidine pathway. The target of olorofim, DHODH, is highlighted. Blue is more than 1-fold downregulated, yellow more than 1-fold upregulated; red is more than 5-fold upregulated. The right of each box is associated with Δ*acdX*, left with Δ*devR*.

Our transcriptomic data and the phenotype of the null mutants led us to assess the effect of pyrimidine pathway precursors on olorofim susceptibility in the transcription factor null mutants. *A. fumigatus* will utilise glutamine as a preferential nitrogen source, even in the presence of other nitrogen containing compounds such as nitrate as pathways that process these precursors are repressed [43, 44]. Intriguingely however, when nitrate was added to the glutamine containing RPMI-1640, the sensitivity of *A. fumigatus* to olorofim increased indicating that even in the presence of preferential nitrogen sources, nitrate can initiate an adaptive response (**Supplemental Figure 4)**. In the olorofim resistant, nitrate non-utilising strain Δ*areA,* addition of nitrate to RPMI reduced susceptibility levels back to that seen for the wild-type. For the Δ*devR* isolate, where the nitrate assimilation pathway as well as all other pathways leading to pyrimidine biosynthesis are upregulated, addition of nitrate did not reduce olorofim susceptibility. The olorofim hypersensitive *acdX* null was the most impacted by changes in nitrogen sources, and counter-intuitively given the downregulation of the NAP in this strain, by the addition of nitrate reduced olorofim susceptibility. These data, combined with results from our transcriptomic analysis suggest that modification of environmental nitrogen sources and or dysregulation of nitrogen metabolism directly impacts changes in olorofim sensitivity.

### Azole mediated antagonism of olorofim is linked to dysregulation of pyrimidine precursor pathways but is not mediated by transcription factors that govern drug resistance

Next, we assessed if the transcription factor null mutants with differential susceptibility to olorofim retained antagonism by voriconazole. To our surprise, antagonism was not affected in these mutants (**Figure 6a, Supplemental Figure 5**). This indicates that antagonism is more complex and potentially requires multiple regulatory factors. This led us to hypothesise that we could affect antagonism by unlinking the pyrimidine pathway from the transcriptional effect of the addition of sub-MIC concentrations of azole. Therefore, we replaced the promoters of *glnA* (AFUB_070010), *pyrABCN* (AFUB_077330) and its paralogues AFUB_025880, *pyrD* (AFUB_085720) and *pyrE* (AFUB_026780) with the doxycycline-regulatable promoter (tetOFF). As expected, replacing the native promoter of *pyrE* with the highly expressing and inducible tetOFF promoter (**Supplemental Figure 6a**), susceptibility to olorofim reduced dramatically when assessed by broth microdilution (**Figure 6b**). In keeping with our hypothesis that genes upstream of *pyrE* are also important in mediating olorofim susceptibility, modest but reproducible decreases in susceptibility were also observed when the promoters of either *pyrABCN* or *pyrD* were replaced. Next, we assessed susceptibility of these mutants on solid medium using a disk assay. Strikingly under the same conditions, susceptibility of the strains to the azoles increased, suggesting that if resistance to olorofim is induced by upregulation of this pathway, strains may well be hypersensitive to the azoles **(Figure 6c)**.

**Figure 6:**
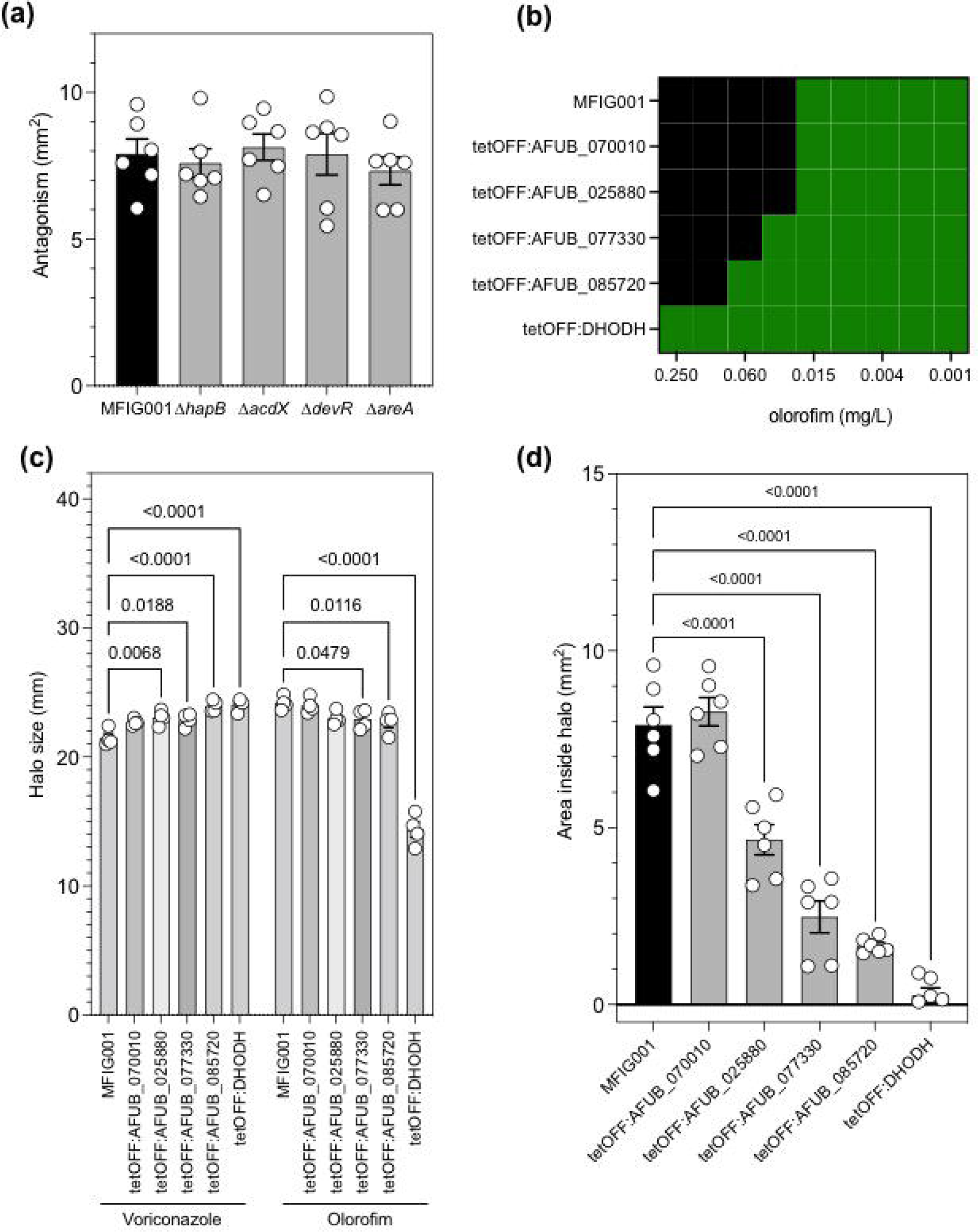
Antagonism between olorofim and the azoles through dysregulation of the pyrimidine pathway. (a) Antagonism for the TF mutants with differential susceptibility to olorofim (n=6) (b) Microbroth dilution assay by EUCAST methodology to olorofim for MFIG001 and the generated tetOFF mutants in the pyrimidine pathway (n=3). (c) The halo size for the generated tetOFF mutants in the pyrimidine pathways to voriconazole and olorofim (n=4). Statistical significance was assessed using a one-way ANOVA with Dunnett’s correction (p<0.05 are shown). (d) Antagonism for the tetOFF mutants within the pyrimidine pathway (n=6). Statistical significance was assessed using a one-way ANOVA with Dunnett’s correction (p<0.05 are shown).

To ensure there was no significant impact on changing the susceptibility of the azoles in our assessment of antagonism in our plate assay, doxycycline levels were titrated to ensure the halo induced by olorofim and voriconazole was almost identical to that of MFIG001 (**Supplemental Figure 6b**). Consistent with our hypothesis that azole induced antagonism is mediated by the pyrimidine biosynthesis pathway antagonism was reduced in a step-wise manner within genes of the pyrimidine pathway, and completely ablated in the tetOFF:DHODH, regardless of the amount of doxycycline used (**Figure 6d, Supplemental Figure 6c**).

In conclusion, we have identified a high-level coordination of the regulation of azole and orotomide resistance, seemingly caused by a crosstalk between the control of the ergosterol and pyrimidine biosynthetic pathways. These pathways are induced in the presence of the azoles resulting in an antagonistic effect on the novel DHODH inhibitor olorofim.

## Discussion

Olorofim is a novel antifungal, currently in phase 3 clinical trials. It has a broad spectrum of activity against most moulds and acts by inhibiting the pyrimidine biosynthetic pathway through disruption of DHODH activity [7]. Our preliminary analysis of the inhibitory effects of olorofim revealed that the MIC and the IC50 were separated over a relatively large concentration range (5-fold). This contrasts with what is seen with itraconazole and other azoles where this concentration spread is typically 2-fold. The clinical implication of this finding remains unclear, however it is likely that olorofim will support clearance of an infection at doses well below the MIC. At these lower concentrations however, exposure to drug will be imparting selective pressure and has the potential to induce the production of mutagenic precursors that may drive the emergence of resistance as has been shown for several antibiotics [45]. As with other anti-infectives that act by inhibiting a single biological target there is clear potential for emergence of resistance. Understanding these mechanisms will provide a framework for development of diagnostics to detect resistance rapidly in the clinic.

Our previous survey of itraconazole sensitivity in the *A. fumigatus* COFUN transcription factor knockout library [46] revealed 6 null mutants that had decreased sensitivity (ranging from 4 to 6-fold increase in MIC compared to the isogenic control) and 6 had increased sensitivity (4 to 8-fold decrease in MIC) to itraconazole. Here our screen revealed that only 1 mutant (Δ*acdX*) showed increased sensitivity while 3 showed decreased sensitivity (Δ*hapB*, Δ*devR*, Δ*areA*) to olorofim and the changes in sensitivity in these isolates were less extreme than seen for the azoles indicating that the frequency of olorofim resistance maybe lower than that seen for itraconazole. Indeed this hypothesis is supported by a recent study that revealed the frequency of olorofim resistance is variable between strain ranging from 1.3 × 10^-7^– 6.9 × 10^-9^, while for itraconazole resistance occurs at an order of magnitude higher (1.2 × 10^-6^ and 3.3 × 10^-8^) [22]. It is unsurprising, given the mechanism of action of olorofim, that the transcription factors that we have identified in this screen either have well defined roles in regulating nitrogen utilisation or have been linked to this function in our study.

What is remarkable however given the distinct mechanisms of actions of the two compound classes, loss of function of either of AreA and HapB results in cross-class resistance to both the azoles and orotomides. HapB is a member of the heterotrimeric CCAAT-binding complex (CBC) and alongside HapC and HapE regulates the expression of over a third of the genome [47] including several genes involved in ergosterol biosynthesis. The *hapB* null displayed the highest levels of resistance to olorofim and was able to germinate at 0.12 mg/L, which is 8-fold higher than the parental isolate but within the concentration range needed for clinical utility. In *A. nidulans* AreA is a positive regulator of many genes that are required for utilisation of nitrogen sources other than glutamate or ammonia [48] with loss of function resulting in an inability to utilise amongst other nitrogen sources, nitrate, nitrite, uric acid and many amino acids [49]. Reassuringly however, drug concentrations in animal models are tolerated well above the increased MIC levels of the null mutants identified in this screen. Dosing 8 mg/kg at 8 hour intervals in mice results in peak serum levels of 2.5-3 mg/L [50]. Olorofim can be tolerated at doses as high as 30 mg/kg intravenously, giving scope for higher drug levels *in vivo* if required. In cynomolgus monkeys a single oral dose of olorofim resulted in peak levels of 0.605-0.914 mg/L in serum for female and male animals, respectively [51].

Our studies have shown there is a clear uni-directional antagonism of the azoles on olorofim, mediated by azole induced overexpression of the pyrimidine biosynthetic pathway and/or metabolic flux through this pathway. Whilst concerning, the antagonism is only evident when relatively low levels of both drugs are used. It is interesting to note that the TR_34_ L98H isolate used in this study has reduced susceptibility to olorofim when compared to the CEA10 isolate and the antagonism drives the MIC above 0.5 mg/L, whether this is of clinical significance remains to be determined. Interestingly, over-expression of any part of the pyrimidine biosynthetic pathway results in a modest increase in susceptibility of *A. fumigatus* to the azoles indicating that some strains that are resistant to olorofim may be more susceptible to the azoles and highlighting that there is complex crosstalk between the ergosterol and pyrimidine biosynthetic pathways. If these drugs are to be used in combination in a clinical setting, careful evaluation of respective drug levels at the site of infection to ensure sufficient concentration of drug to avoid antagonism would be sensible. The consequences of using azoles and olorofim in combination for treatment of strains harbouring the TR_34_ L98H allele also needs further evaluation.

In summary, we have explored the mechanism behind olorofim susceptibility through a systematic analysis of the COFUN transcription factor null library. All the mutants we identified that had altered sensitivity to olorofim have associated defects in nitrogen metabolism. Two of these mutants, Δ*devR* and Δ*acdX*, show dysregulation of genes involved in metabolic pathways immediately upstream of the pyrimidine pathway potentially leading to a differential flux of metabolites into this pathway. Importantly, we have identified two transcription factors, the CBC and AreA, that regulate cross resistance to both the azoles and olorofim. Lastly, we have detected an antagonistic effect between olorofim and the azoles which we can modulate through transcriptionally unlinking the pyrimidine pathway from upstream pathways.

## Acknowledgements

The authors would like to thank F2G for supplying the antifungal olorofim. We would also like to thank the Genomic Technology Core Facility, the Bioinformatics Core Facility in the University of Manchester for their technical support. This work was supported by the Wellcome Trust grant number 219551/Z/19/Z and 208396/Z/17/Z to M.B. CV is funded by postdoctoral fellowship from Fundação de Amparo à Pesquisa do Estado de São Paulo (FAPESP-BEPE 2020/01131-5).

## Author contributions

N.v.R designed and performed the experiments, analysis, wrote and edited the manuscript. S.H. designed and performed experiments and analysis. I.S. designed and performed experiments and analysis. C.V. designed and performed experiments and analysis. H.A. designed and performed experiments and analysis. G.G. provided funding and edited the manuscript. F.G. designed and performed experiments and analysis. J.A. designed and performed experiments and analysis. M.B. provided funding, designed the experiments, wrote and edited the manuscript

---

**Supplemental Figure 1:**
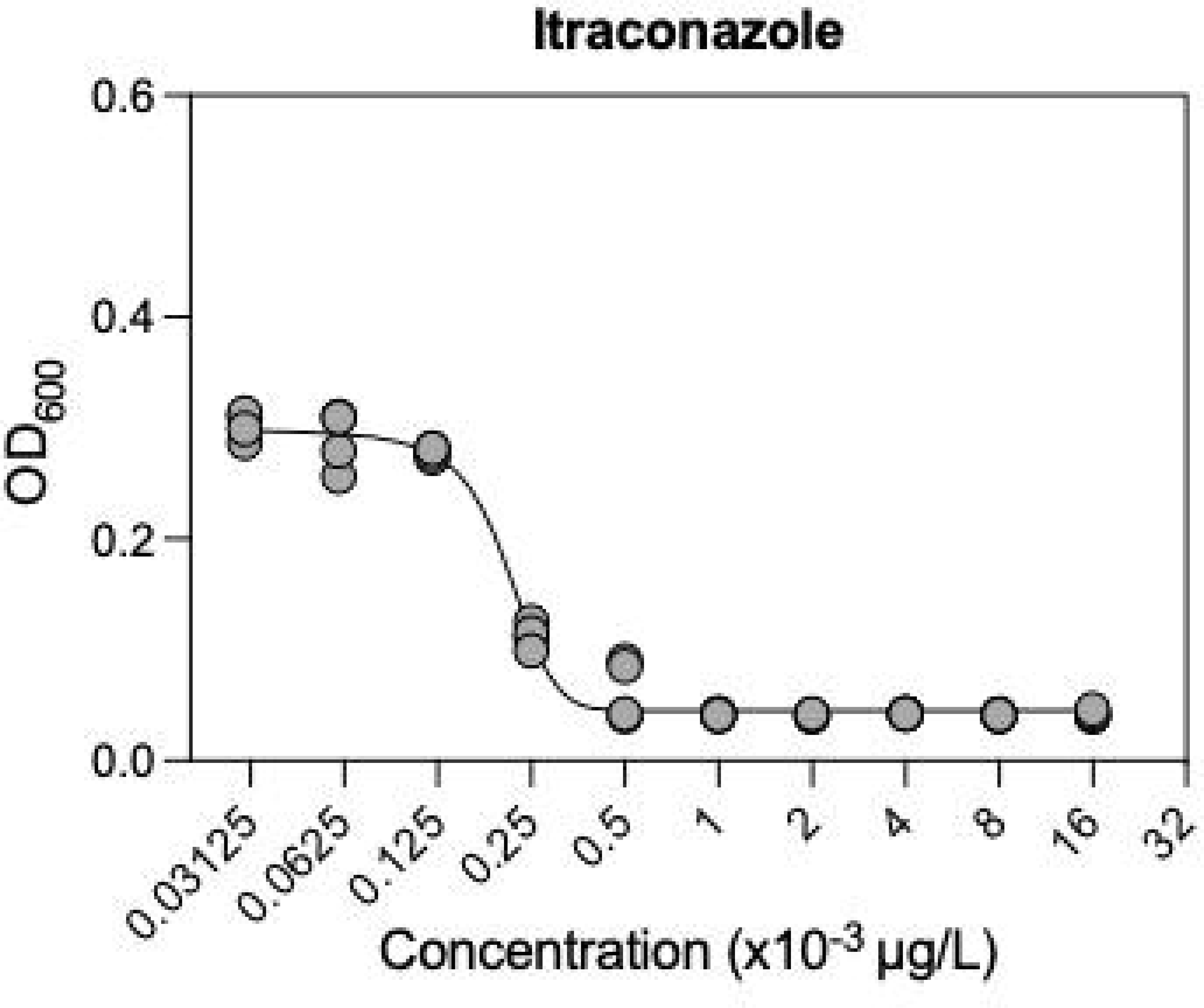
Determination of IC50 for itraconazole. MIC to olorofim in RPMI-1640 was determined according to EUCAST methodology for *A. fumigatus* MFIG001. OD600 was measured after 48 hours to determine growth quantitatively.

**Supplemental Figure 2:**
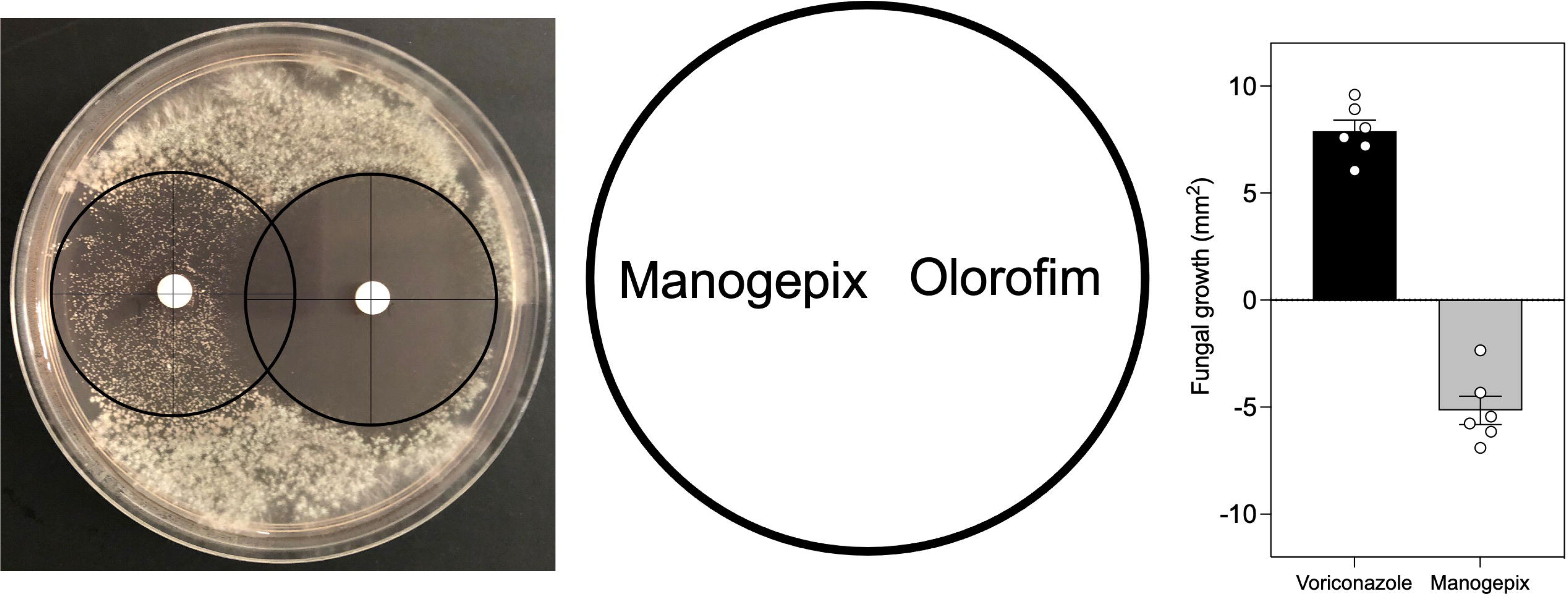
Antagonism between manogepix and olorofim. Antagonism was assessed and quantified (n=6) between manogepix and olorofim. A synergy between these two novel antifungals was observed.

**Supplemental Figure 3:**
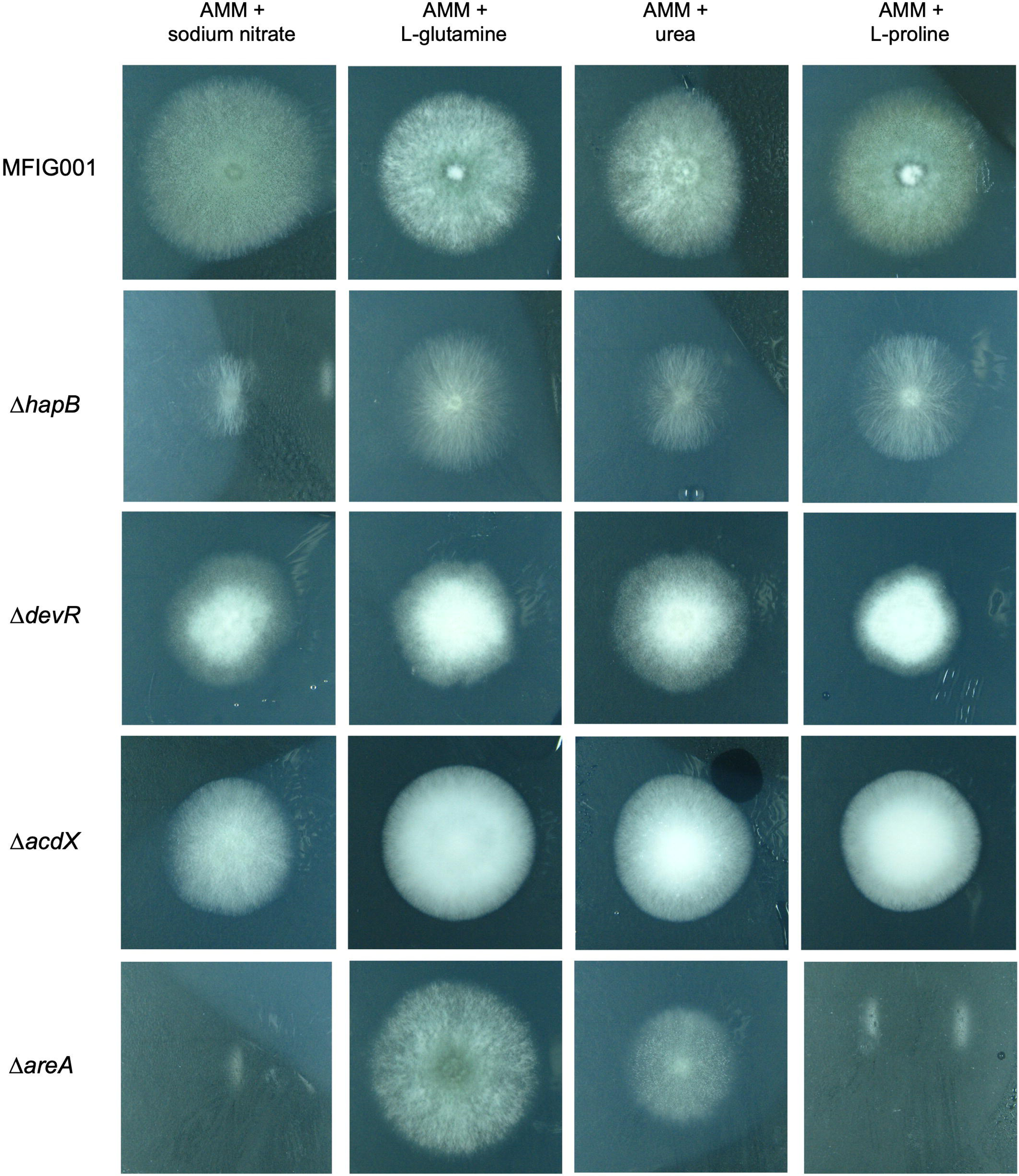
Images of nitrogen spot tests of TF null mutants. Growth of TF null mutants and wild-type was assessed on AMM supplemented with 50 mM ammonium tartrate, 10 mM sodium nitrate, 10 mM L-glutamine, 10 mM urea or 10 mM L-proline (n=3). Images were taken after 72 hours at 37 Celsius.

**Supplemental Figure 4:**
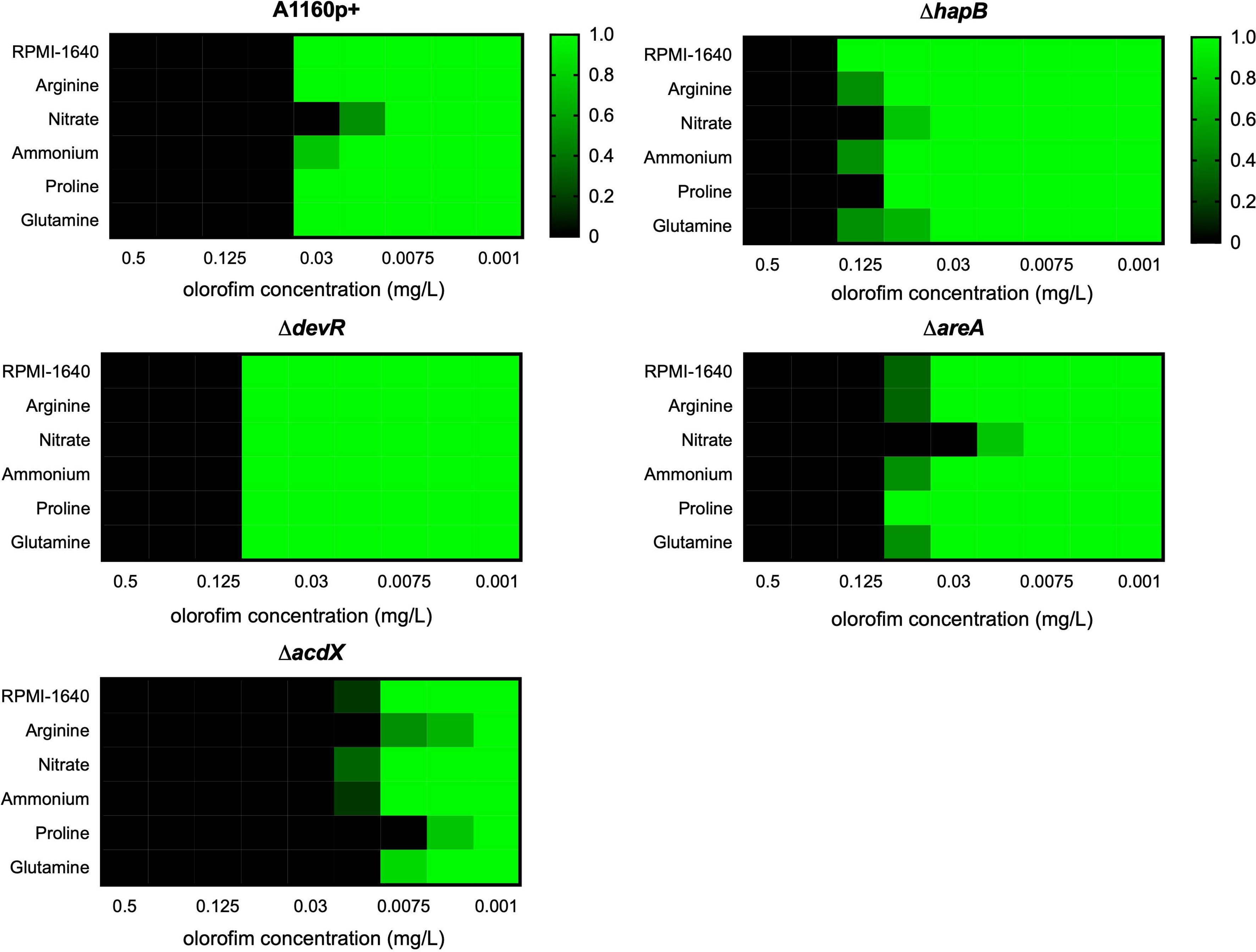
The effect of nitrogen source on olorofim susceptibility. MICs according to EUCAST methodology in RPMI-1640 supplemented with either 20 mM arginine, 10 mM nitrate, 20 mM proline or 50 mM glutamine. Addition of nitrate changed Olorofim susceptibility by 2-fold for all strains except Δ*devR*.

**Supplemental Figure 5:**
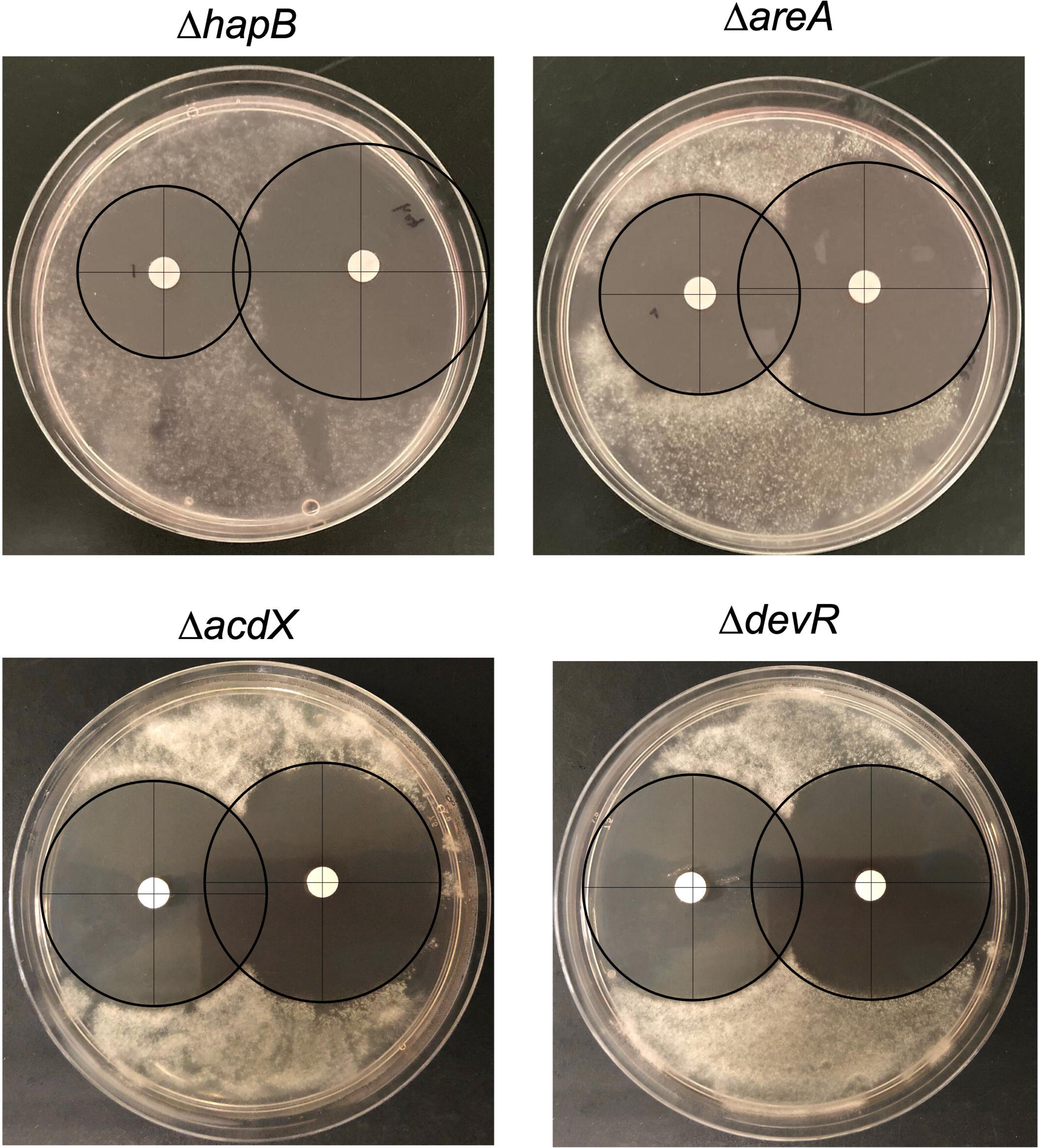
Antagonism of the azoles to olorofim for TF mutants. Representative images of TF mutants with differential susceptibility to olorofim tested for antagonism between voriconazole (800 mg/L) and olorofim (500 mg/L).

**Supplemental Figure 6:**
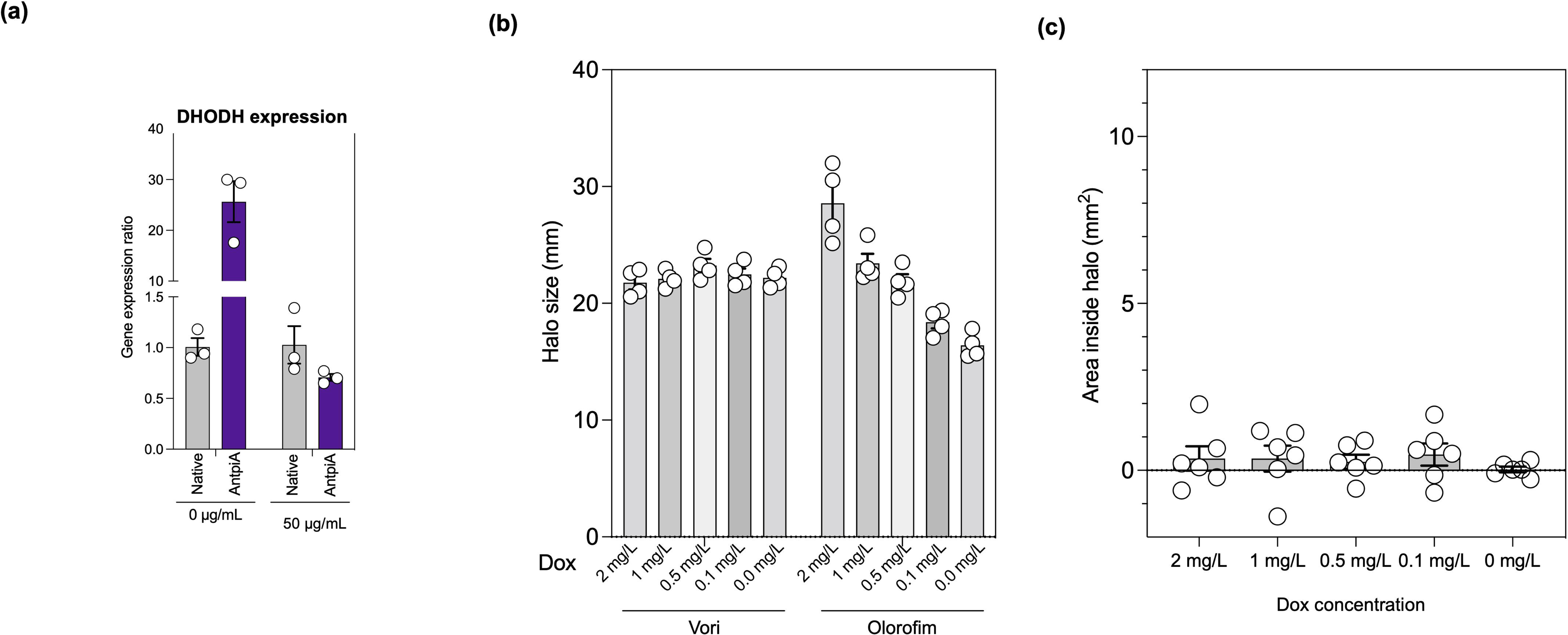
Overexpression of DHODH by inserting the tetOFF cassette. **(a)** The DHODH gene was overexpressed by inserting the tetOFF cassette as a promoter system. Expression was measured by qPCR (n=3) without doxycycline and could be reduced by addition of doxycycline (50 µg/mL). (b) The halo size for tetOFF:DHODH at different doxycycline concentrations. (c) The area inside the halo at different concentrations of doxycycline (n=6) for the tetOFF:DHODH mutant.

**Supplemental Table 1: Oligos and crRNA used in this study.**

